# Differential effects of antiseptic mouth rinses on SARS-CoV-2 infectivity in vitro

**DOI:** 10.1101/2020.12.01.405662

**Authors:** Chuan Xu, Annie Wang, Eileen R. Hoskin, Carla Cugini, Kenneth Markowitz, Theresa L. Chang, Daniel H. Fine

## Abstract

SARS-CoV-2 is detectable in saliva from asymptomatic individuals, suggesting a potential benefit from the use of mouth rinses to suppress viral load and reduce virus spread. Published studies on reduction of SARS-CoV-2-induced cytotoxic effects by antiseptics do not exclude antiseptic-associated cytotoxicity. Here, we determined the effect of commercially available mouth rinses and antiseptic povidone-iodine on the infectivity of SARS-CoV-2 virus and of a non-pathogenic, recombinant, SARS-CoV-2 infection vector (pseudotyped SARS-CoV-2 virus). We first determined the effect of mouth rinses on cell viability to ensure that antiviral activity was not a consequence of mouth rinse-induced cytotoxicity. Colgate Peroxyl (hydrogen peroxide) exhibited the most cytotoxicity, followed by povidone-iodine, chlorhexidine gluconate (CHG), and Listerine (essential oils and alcohol). Potent anti-viral activities of povidone iodine and Colgate peroxyl mouth rinses was the consequence of rinse-mediated cellular damage. The potency of CHG was greater when the product was not washed off after virus attachment, suggesting that the prolonged effect of mouth rinses on cells impacts anti-viral activity. To minimalize mouth rinse-associated cytotoxicity, mouth rinse was largely removed from treated-viruses by centrifugation prior to infection of cells. A 5% (v/v) dilution of Colgate Peroxyl or povidone-iodine completely blocked viral infectivity. A similar 5% (v/v) dilution of Listerine or CHG had a moderate suppressive effect on the virus, but a 50% (v/v) dilution of Listerine or CHG blocked viral infectivity completely. Prolonged incubation of virus with mouth rinses was not required for viral inactivation. Our results indicate that mouth rinses can significantly reduce virus infectivity, suggesting a potential benefit for reducing SARS-CoV-2 spread.

**Importance:** SARS-CoV-2 is detectable in saliva from asymptomatic individuals, suggesting the potential necessity for the use of mouth rinses to suppress viral load to reduce virus spread. Published studies on anti-SARS-CoV-2 activities of antiseptics determined by virus-induced cytotoxic effects cannot exclude antiseptic-associated cytotoxicity. We found that all mouth rinses tested inactivated SARS-CoV-2 viruses. Listerine and CHG were less cytotoxic than Colgate Peroxyl or povidone-iodine and were active against the virus. When mouth rinses were present in the cell culture during the infection, the potent anti-viral effect of mouth rinses were in part due to the mouth rinse-associated cytotoxicity. Our results suggest that assessing anti-viral candidates including mouth rinses with minimal potential disruption of cells may help identify active agents that can reduce SARS-CoV-2 spread.

## Introduction

Severe acute respiratory syndrome-related coronavirus (SARS-CoV-2), a non-segmented positive-strand RNA enveloped virus, is the causative agent of coronavirus disease 19 (COVID-19). As of November 25, 2020, **COVID**-19 spread has resulted in more than 59 million cases worldwide (https://covid19.who.int/) and more than 12.3 million cases in the United States alone (https://covid.cdc.gov/covid-data-tracker/#cases_casesper100klast7days). Evidence indicates that transmission of SARS-CoV-2 occurs through virus-containing secretions, such as saliva and respiratory secretions, or their droplets (WHO 2020). Salivary SARS-CoV-2 viral load is highest during the first week after symptom onset (To et al. 2020), and individuals with SARS-CoV-2 infection shed virus and can remain asymptomatic for a prolonged period (Lee et al. 2020; Wei et al. 2020), highlighting the importance of developing a strategy to prevent virus spread in the general population. Additionally, there is an urgent need for evidence-based practices to protect patients and healthcare workers in the dental office and elsewhere when salivary droplets and aerosols are generated during dental treatment when masks for patients are not an option.

Antiseptic mouth rinses have been shown to have efficacy in reducing bacteria and viruses in the oral cavity and in dental aerosols (Fine et al. 1993; Fine et al. 1996; Koletsi et al. 2020). The antiseptic Listerine and chlorhexidine gluconate-0.12% (CHG) have been shown to reduce herpes simplex virus-1 load in saliva after rinsing (Meiller et al. 2005; Park and Park 1989). Potential inhibitory effects of mouth rinses on SARS-CoV-2 inactivation have been proposed based on the assumption that the organic components in the mouth rinses disrupt viral membranes (O’Donnell et al. 2020). Because viable cells are required for productive infection, toxic effects on cells that may produce unfavorable conditions for viral infection can be misinterpreted as a potent antiviral activity. This has been an issue in recent studies on the effect of mouth rinses and povidone-iodine on SARS-CoV-2 infection in which mouth rinse-associated cytotoxic effects were not excluded (Anderson et al. 2020; Bidra et al. 2020; Meister et al. 2020). Thus, interpretation of anti-viral effects of mouth rinse is complicated because the anti-viral effects may be the consequence of cytotoxicity. For example, in the study by Bidra et al, the mixture of SARS-CoV2 viruses and diluted povidone-iodine was added to Vero cells for 5 days followed by a determination of cytopathic effects. The same assay was conducted by Meister et al, wherein cell viability was determined by crystal violet staining, a method that does not directly distinguish live and dead cells. Antiseptic-associated cell death can result in decreased numbers of target cells for viral infection producing an apparent decrease in viral infectivity, which can be mistaken as a potent anti-viral effect.

Here, we determined the effect of mouth rinses and antiseptics including Listerine, chlorhexidine gluconate (CHG), povidone-iodine, Colgate Peroxyl on cell viability prior to assessing their impact on the infectivity of SARS-CoV-2 viruses. We used replication competent SARS-CoV-2 viruses expressing mNeonGreen, which allowed us to monitor the green signal in live cells within 24 h after infection which avoided significant virus-induced cytopathic effects seen at later time points. We also employed a single-cycle infection assay using a pseudotyped SARS-CoV-2 virus expressing SARS-CoV-2 spike proteins, which provides a non-pathogenic vector for assessing viral infectivity. Pseudotyped virus does not cause virus-induced cytotoxic effects, but allows us to assess SARS-CoV-2 spike protein-mediated viral entry. We tested the effects of serial dilutions of the mouth rinses to determine their relative effectiveness against the virus directly as opposed to their cytotoxicity against mammalian cells. Overall, Listerine and CHG were less cytotoxic than Colgate Peroxyl or povidone-iodine, and were active against the virus. When mouth rinses were present in the cell culture during the infection, the anti-viral effect of mouth rinses appeared to be more potent, an apparent consequence of mouth rinse-associated cytotoxicity. Our results suggest that assessing anti-viral candidates, including mouth rinses, under conditions of minimal potential disruption of cells will help identify active agents that can reduce SARS-CoV-2 spread.

## Materials and Methods

### Reagents

The infectious-clone-derived SARS-CoV-2 virus (USA_WA1/2020 strain) expressing mNeonGreen was kindly provided by Pei-Yong Shi at the University of Texas Medical Branch, Galveston, TX USA (Xie et al. 2020). A recombinant construct used for infectivity assays (pseudotyped SARS-CoV-2) was derived from the full-length SARS-CoV2-Wuhan-Hu-1 surface (spike) (GenBank accession number QHD43416)(Wu et al. 2020), which was codon optimized for humans and synthesized with Kozak-START GCCACC ATG and STOP codons, flanked by 5’ Nhel/3’Apal sites for subcloning into the pcDNA3.1(+) vector (Thermo Fisher Scientific, USA). HEK293T cells and Vero E6 cells were purchased from ATCC. Monoclonal antibody (Ab) against SARS-CoV-2 spike protein (IgG1 clone#43, Cat # 40591-MM43) was purchased from Sino Biological, Inc (Wanye, PA). Listerine Original (Johnson & Johnson Consumer Inc, Skillman, NJ, USA), povidone-iodine-10% (1% available iodine, CVS Pharmacy Inc, Woonsocket, RI, USA), Colgate Peroxyl (1.5% w/v hydrogen peroxide, Colgate-Palmolive Inc, New York, NY, USA), and Chlorhexidine Gluconate-0.12% (Xttrium Laboratories Inc, Mount Prospect, IL, USA) (Table 1) were purchased from a local pharmacy.

**Table 1.** Mouth rinse and antiseptic products used in this study

| **Rinse** | **Manufacturer** | **Active ingredients** |
| --- | --- | --- |
| Listerine Antiseptic Original | Johnson & Johnson Consumer Inc, Skillman, NJ, USA | 20-30% Ethanol<br>Thymol 0.064%<br>Methyl salicylate 0.06%<br>Menthol (Racemethol) 0.042%<br>Eucalyptol 0.092% |
| Povidone-Iodine | CVS Pharmacy Inc, Woonsocket, RI, USA | 10% solution<br>(1% available iodine) |
| Colgate Peroxyl | Colgate-Palmolive Inc, New York, NY, USA | 1.5% w/v hydrogen peroxide |
| Chlorhexidine Gluconate | Xttrium Laboratories Inc, Mount Prospect, IL, USA | 0.12% Chlorhexidine |

### Cell culture

HEK293T cells and human angiotensin-converting enzyme 2 (hACE2)-expressing HeLa cells (kindly provided by Dennis Burton; The Scripps Research Institute, La Jolla, CA) were cultured in Dulbecco’s Modified Eagle’s Medium (DMEM) supplemented with 10% fetal bovine serum (FBS). Vero E6 cells were cultured in Eagle’s Essential Minimal Medium (EMEM) with 5% FBS. TR146 cells were cultured in Ham’s F12 media with glutamate and 10% FBS.

### Viral infection

Replication competent SARS-CoV-2 viruses expressing mNeonGreen were propagated in Vero E6 cells as described previously (Xie et al. 2020). All experiments were performed in a biosafety level 3 (BSL3) laboratory. Powered air purifying respirators (Breathe Easy, 3M), Tyvek suits, aprons, sleeves, booties, and double gloves were worn. Virus titers were determined by plaque assays. Briefly, Vero E6 cells were seeded at 6×10^5^ cell per well in a 6-well plate and cultured for overnight. Cells were then exposed to serial dilutions of SARS-CoV-2 viruses for 1.5-2 h. After removing unbound viruses, cells were overlayed with 0.8% Agarose LE (Sigma) in DMEM with 2%FBS. On post-infection day 3, cells were fixed with 10% formaldehyde (in PBS) for 30 min. Agarose plugs were removed, and fixed cells were stained with 0.2% crystal violet (w/v) in ethanol.

For the infection assay, Vero E6 cells at 1×10^4^ cells per well were incubated overnight in a black 96-well glass plate (Greiner). Cells were exposed to treated or untreated viruses in 50 μl at a multiplicity of infection (MOI) of 5 for 1 h followed by the addition of 100 μl FluoroBrite medium containing 2% FBS. The fluorescent signal from productive viral infection and cell images were monitored at 24 h after infection by using a Biotek Cytation 5.

For single-cycle infection assay, replication-defective HIV-1 luciferase-expressing reporter viruses pseudotyped with SARS-CoV2 S proteins were produced by co-transfection of a plasmid encoding the envelope-deficient HIV-1 NL4-3 virus with the luciferase reporter gene (pNL4-3.Luc.R+ E-, kindly provided by Nathaniel Landau, New York University) and a pcDNA3.1 plasmid expressing the SARS-CoV2 glycoprotein into HEK 293T cells using Lipofectamine 3000 (Thermo Fisher Scientific). The supernatant was collected 48 h after transfection, and filtered. Virus stocks were analyzed for HIV-1 p24 antigen by the AlphaLISA HIV p24 kit (PerkinElmer). Virus stocks contained approximate 200 ng/ml of HIV p24 protein.

For infection assays, cells were seeded at 5×10^4^ cells/well in a 48-well plate and cultured overnight. Pseudotyped SARS-CoV2 luciferase reporter viruses were incubated with or without mouth rinse for 30 min at 37°C before being added to HeLa-hACE2 cells. After 1-2 h viral attachment, infected cells were cultured in media with 10% FBS for 48-72 h. Cells were then lysed in 1x passive lysis buffer (Promega Inc.) followed by measuring luciferase activity (relative light units; RLUs) using Luciferase Substrate Buffer (Promega Inc) on a 2300 EnSpire Multilabel Plate Reader (PerkinElmer, Waltham, MA).

To assess the effect of mouth rinses on the viruses, mouth rinse-treated SARS-CoV-2 viruses were concentrated by centrifugation at 14,000 rpm in a centrifuge (Eppendorf) at 4°C for 2 h as described previously (Holmes et al. 2015). After removing the mouth rinses or media (control samples), virus pellets were resuspended in DMEM and used to infect HeLa-hACE2 cells. Infection was determined by measuring fluorescence intensity after 25 h for replication competent viruses or luciferase activity after 48 h for pseudotyped viruses.

### Cytotoxicity Assay

HeLa-hACE2 and TR146 cells were plated in 96-well plates at 5,000 cells per well, and then treated with various dilutions of mouth rinses for the times indicated in the figure legends. Cell viability was analyzed using CellTiter 96® AQueous One Solution Cell Proliferation Assay (Promega, Madison, WI) according to the manufacture’s instruction. The reagent contains a tetrazolium compound [3-(4,5-dimethylthiazol-2-yl)-5-(3-carboxymethoxyphenyl)-2-(4-sulfophenyl)-2H-tetrazolium, inner salt; MTS] and an electron coupling reagent (phenazine ethosulfate; PES) to measure metabolically active cells.

### Statistical analysis

Statistical comparisons were performed using one-way ANOVA Dunnett’s multiple comparisons test or two-tailed Mann-Whitney U test as appropriate. Prism 8 (GraphPad Software, LLC) was used. *p* < 0.05 was considered significant.

## Results

### Differential effects of mouth rinses on cell viability

It is critical to assess antiviral agents under non-cytotoxic conditions as viruses depend on viable host cells for productive infection. Therefore, we first determined the effect of mouth rinses on cell viability. Note that percentage dilutions (v/v) of commercial mouth rinse products are referenced in this study. For example, in Figure 1, 50% (v/v) CHG represents a solution composed of equal volumes of culture media and of the commercial product, and does not indicate the final concentration of active ingredients. HeLa-hACE2 cells were treated with 2-fold serial dilutions in medium of Listerine, CHG, povidone-iodine, or Colgate Peroxyl for 20 sec, washed, cultured in fresh media, and cell viability was determined. All 50% (v/v) dilutions of mouth rinses were highly toxic to HeLa-hACE2 and oral epithelial cells (**Fig 1**). Listerine was least cytotoxic, followed closely by CHG. Both 0.5% (v/v) dilutions of povidone-iodine and Colgate Peroxyl were highly toxic to cells.

**Fig. 1.**
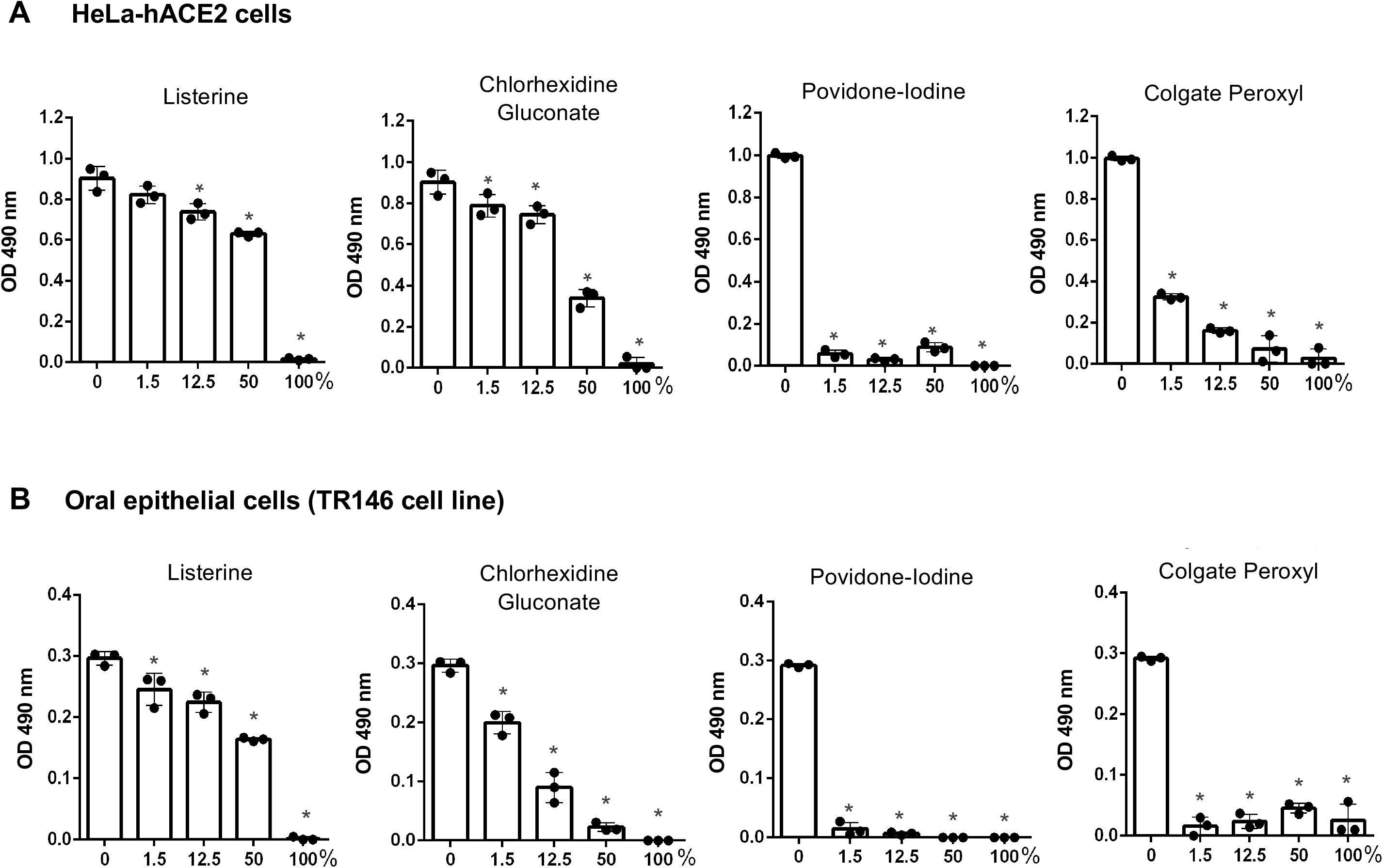
The effect of short-term exposure of mouth rinses on the viability of HeLa-hACE2 and oral epithelial cells. hACE2-expressing HeLa cells (A) and oral epithelial TR146 cells (B) were treated for 20 sec with different dilutions (v/v) of products including Listerine, CHG, povidone-iodine, or Colgate Peroxyl. Cells were washed and cultured with fresh media immediately. Cell viability was determined by MTS-based CellTiter 96® AQueous One Solution Cell Proliferation Assay. Data are mean ± SD of 3 samples. Significance of differences between mouth rinse-treated cells and mocked-treated controls was compared; \**p* < 0.05.

We also determined the effect of 2h exposure of mouth rinses on cell viability for comparison with the duration of viral attachment in the infection assay. We found that 6.25% (v/v) diluted Listerine and 1.5% (v/v) diluted CHG did not impact cell viability, whereas 0.1% (v/v) diluted povidone-iodine or Colgate Peroxyl significantly affected cell viability after 2 h exposure (**Fig 2**).

**Fig. 2.**
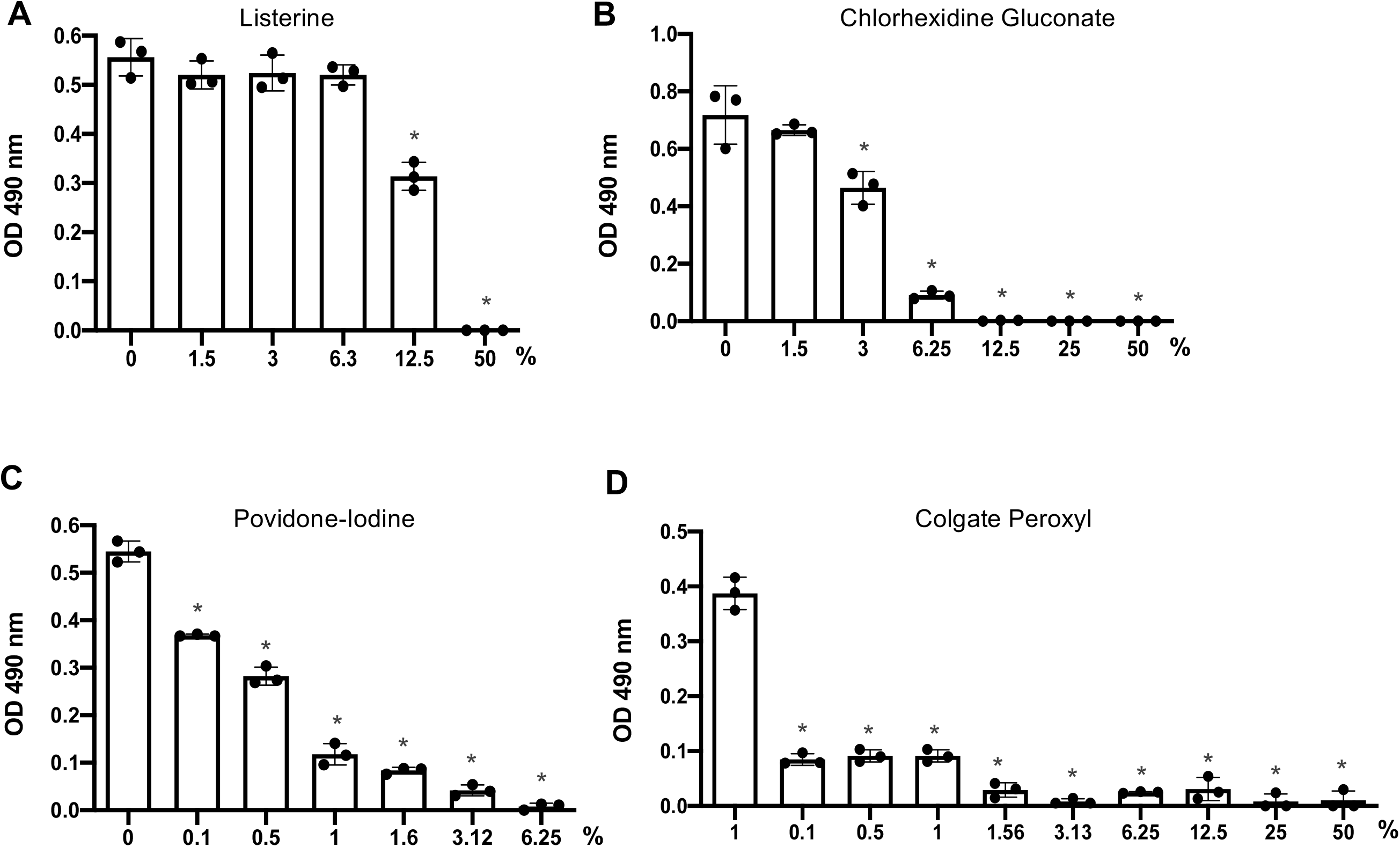
The effect of prolonged exposure to mouth rinses on cell viability. hACE2-expressing HeLa cells were treated for 2 h with serial dilutions of products including Listerine (A), CHG (B), povidone-iodine (C), or Colgate Peroxyl (D) starting at 50% (v/v) except povidone-iodine, which was started at 6.25% (v/v). Cell viability was determined by MTS-based CellTiter 96® AQueous One Solution Cell Proliferation Assay. Data are mean ± SD of 3 samples, and are representative of two independent experiments. Significance of differences between mouth rinse-treated cells and mocked-treated controls was compared; \**p* < 0.05.

### *Antiviral effect of diluted* povidone-iodine or Colgate Peroxl was associated with cytotoxicity

Previous studies on the effect of antiseptics on SARS-CoV-2 infection employed methods involving virus-induced cytopathic effects without excluding mouth rinse-associated cytotoxic effects (Anderson et al. 2020; Bidra et al. 2020; Meister et al. 2020). To examine this more closely, we assessed the effect of highly diluted mouth rinse and povidone-iodine on replication competent SARS-CoV-2 viruses without washing off the antiseptics; cell morphology was monitored as a crude measure of cytopathic effects. Viruses were treated with non-cytotoxic dilutions of Listerine and CHG, low cytotoxic dilutions of povidone-iodine, and highly dilute Colgate Peroxyl (with cytotoxic effects) (**Fig 2**), and were immediately added to Vero cells. Additional media were added 2 h after infection, and cells were cultured overnight. The fluorescence intensity from SARS-CoV-2 infection was determined, and cell morphology was imaged at 24 h after infection. Diluted Listerine (3% v/v)CHG reduced SARS-COV-2 infection by 40%, and CHG (1.5% v/v) reduced infection by 70%, without apparent impacts on cell morphology (**Fig 3**). Diluted povidone-iodine (0.1% v/v) and Colgate Peroxyl (0.05% v/v) appeared to have potent anti-viral activities; however, disruption of cell morphology was apparent (**Fig 3**), indicating that the putative anti-viral effect of these two agents was likely a consequence of cytotoxicity.

**Fig 3.**
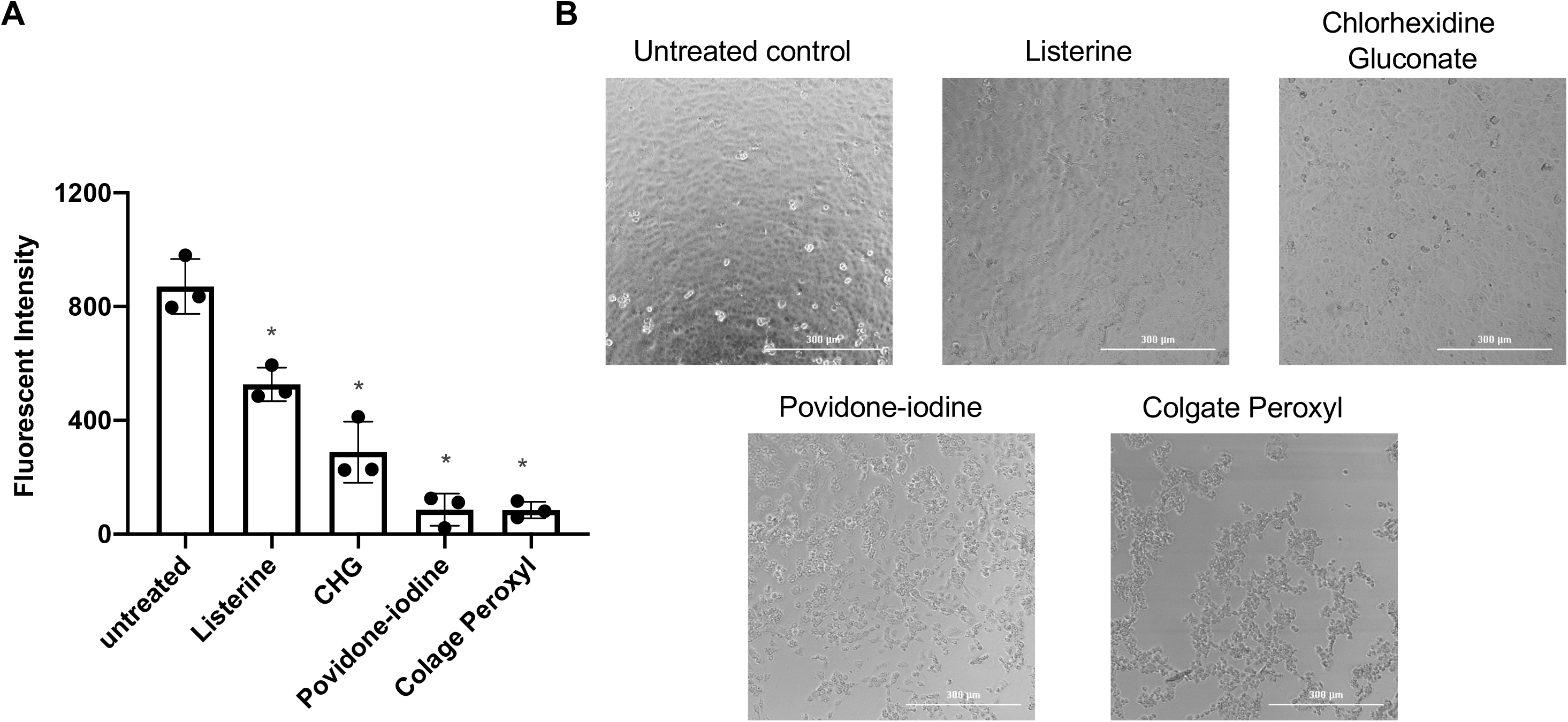
Effect of diluted antiseptics on infection by replication competent SARS-CoV-2 virus when antiseptics were present in the culture. (A) Replication competent virus SARS-CoV-2 expressing mNeonGreen (MOI of 5) were mixed or not with Listerine (3%), CHG (1.5%), povidone-iodine (0.1%), or Colgate Peroxyl (0.05%), and immediately added (in 50 μl) to Vero cells, and incubated 1 h for viral attachment. Antiseptics were not washed off to compare published studies but were diluted by the addition of 100 μl of media to reduce potential toxic effects. Fluorescence intensity derived from productive viral infection was determined at 24 h post infection. Cell images were acquired by using Bioteck Cystatin 5 plate reader. Differences between mouth rinse-treated viruses and medium control (0%) were compared; \**p* < 0.05. Data are means ± SD, and are representative of three independent experiments. CHG

We also used HIV pseudotyped luciferase virus particles expressing SARS-CoV2 surface protein (spike, S) to assess the effect of non-cytotoxic diluted Listerine and CHG on viral infectivity. Unlike replication competent SARS-CoV-2 virus, which induces cytopathic effects after prolonged culture, HIV pseudotyped luciferase viruses provide a reliable, non-pathogenic, vector for assessing viral infectivity. We confirmed that infection by pseudotyped SARS-CoV-2 was dependent on human hACE2, and that infection was neutralized by anti-spike monoclonal antibody (**Supplemental Figure 1**).

We determined the effect of diluted Listerine (1.5-6%v/v) and CHG (1.5% and 3%v/v), which had no or little effect on cell viability (**Fig 2**), on pseudotyped SARS-CoV-2 virus infection without washing off the mouth rinses during the infection. We found that 6% (v/v) Listerine had a moderate anti-SARS-CoV-2 activity, whereas 1.5% or 3% (v/v) CHG suppressed viral infection by 88% and 97%, respectively (**Fig 4A**).

**Fig. 4.**
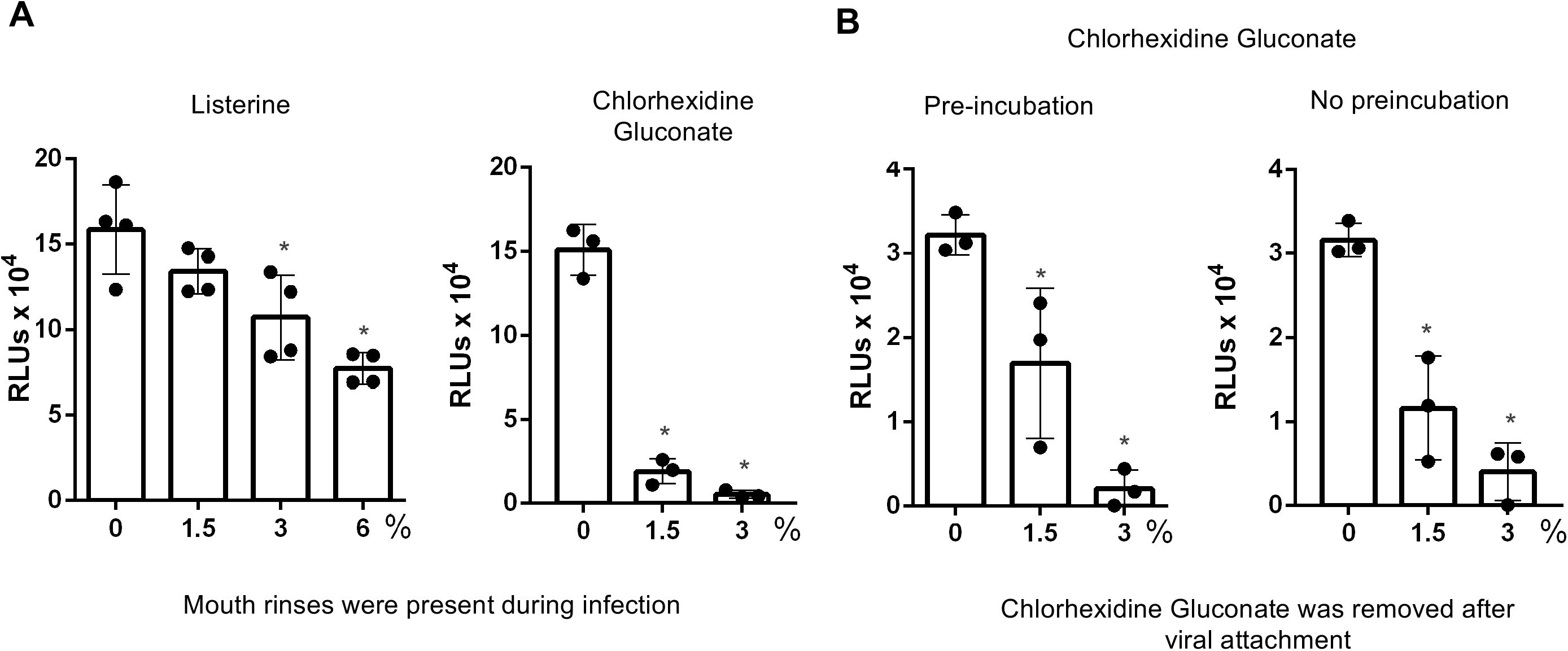
Effect of non-cytotoxic diluted Listerine and Chlorhexidine Gluconate on infection by pseudotyped SARS-CoV-2 virus. (A) Pseudotyped SARS-CoV-2 virus (100 μl) was incubated with or without diluted Listerine or CHG at non-cytotoxic concentrations at 37°C for 30 min, and then added to HeLa-hACE2 cells. After 1-2 h incubation, an additional 400 μl of DMEM 10% FBS was added to the cells without washing off viruses or mouth rinses. Infected cells were cultured in the presence of mouth rinses for 2 days before measuring luciferase activity. (B) Pseudotyped SARS-CoV-2 virus were pre-incubated with diluted CHG at 37°C for 30 min (left panel) or without 30 min preincubation (right panel). Treated viruses were added to HeLa-hACE2 cells for 1 h at 37°C. Cells were washed to remove residual mouth rinse, and cultured for 2 days before measuring luciferase activity. Significance of differences between mouth rinse-treated viruses and mocked-treated controls was compared; \**p* < 0.05. Data are means ± SD.

We also determined whether pre-incubation of viruses with CHG affected the degree of anti-viral activity. For this, pseudotyped SARS-CoV-2 viruses were pre-treated or not with CHG 30 min at 37°C before being added to target cells. In contrast to the experiment shown in **Fig 4A**, in which the mouth rinses were present during infection, here, the mixture of virus and CHG was removed, fresh media were added, and cells were cultured for 2 days before measuring luciferase activity (**Fig 4B**). The effect of CHG with or without pre-incubation was comparable (**Fig 4B**). We found that the anti-viral effect of 1.5% (v/v) CHG was less potent when CHG was present only during viral attachment (**Fig 4B**) compared to being continuously present during viral infection and incubation for 2 days (**Fig 4A**). The more pronounced anti-viral activity of 1.5% (v/v) CHG in this experiment may be due to effects of prolonged contact with CHG on target cells, indicating the importance of minimizing mouth rinse-associated cytotoxicity in the infection assay.

### Direct effect of mouth rinses on viruses

To assess direct effects of the rinses on virus particles, pseudotyped SARS-CoV-2 viruses were incubated with mouth rinses for 30 min at 37°C, and were pelleted by centrifugation (Holmes et al. 2015) prior to the infection assay. Note that centrifugation did not impact infectivity of the virus (data not shown). After removal of the supernatant containing the mouth rinse, viruses were resuspended in media and added to HeLa-hACE2 target cells (**Fig 5A and 5B**). We also assessed the effect of centrifugation and virus resuspension on infectivity cell viability to monitor potential cytotoxic effects of residual mouth rinses on the cells (Fig 5C). The result showed that all antiseptics tested inactivated viruses. Anti-viral activity of 50% (v/v) Colgate Peroxyl was associated with residual mouth rinse-induced cytotoxicity; 5% (v/v) Colgate Peroxyl and 5% (v/v) povidone-iodine blocked viral infectivity (**Fig 5B**); 50% (v/v) Listerine and 50% CHG inactivated viruses, but 5% of these rinses did not (**Fig. 5B**).

**Fig. 5.**
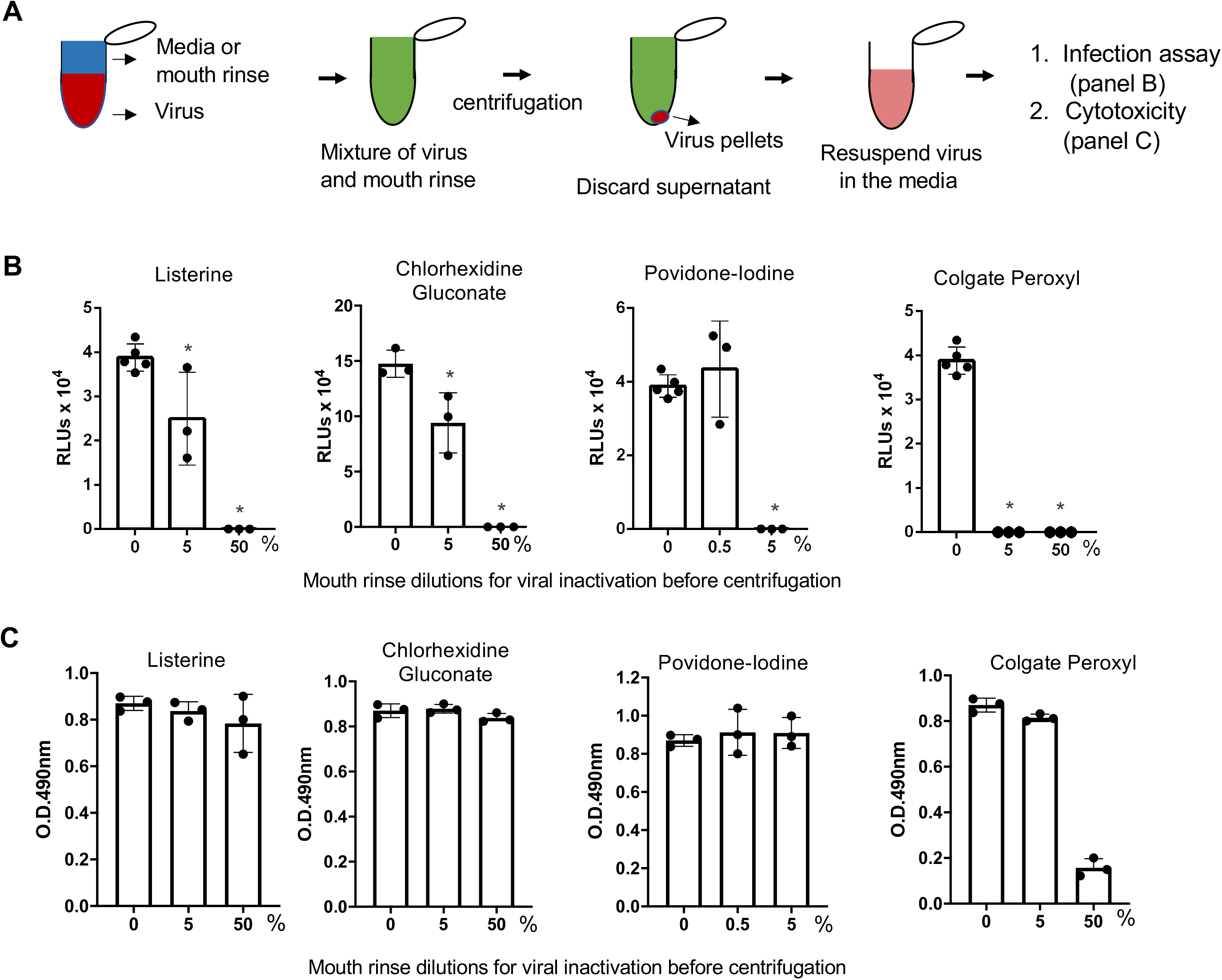
The effect of mouth rinses on SARS-CoV-2 viruses. (A) Experimental design. Pseudotyped luciferase reporter viruses expressing SARS-CoV-2 S proteins were incubated with the indicated dilutions of mouth rinses at 37°C for 30 min followed by centrifugation and aspiration of the supernatant. Virus pellets were resuspended in the culture medium and added to HeLa-hACE2 cells for (B) infection assay and for (C) cytotoxicity assay as described in Methods. Differences between mouth rinse-treated viruses and media controls (0%) were compared; \**p* < 0.05. Data are means ± SD, and are representative of two independent experiments.

Viruses treated with 50% (v/v) Listerine, 50% (v/v) CHG, 50% (v/v)Colgate Peroxyl, or 5% (v/v) povidone-iodine completely lost infectivity (**Fig 5B**); the apparent anti-viral effect of 50% (v/v) Colgate Peroxyl was associated with cell toxicity (**Fig 5C**). Treatment with 5% (v/v) Listerine or CHG had a moderate anti-viral effect; whereas 5% (v/v) Povidone-Iodine or 5% (v/v) Colgate Peroxyl completely inactivated the viruses. Centrifugation and resuspension of the virus had no apparent impact on infectivity. All mouth washes at non-cytotoxic levels exhibited antiviral activity. Colgate Peroxyl and povidone-iodine had greater inhibitory effects on the viruses than CHG or Listerine.

Unlike high concentrations of Colgate Peroxyl and povidone-iodine, whose anti-viral activities were associated with cytotoxicity, higher concentrations of Listerine and CHG exhibited potent anti-viral effects without cytotoxicity. We asked whether preincubation of the virus with Listerine or CHG was required to achieve their direct effect on the virus. Mouth rinses were added to the virus, mixed, and immediately centrifuged at 4°C. Supernatants containing the mouth rinses were discarded. Viruses were then resuspended in media and added to target cells. The viral inhibition profiles of Listerine and CHG without preincubation (**Supplemental Fig 2**) were comparable to those with 30 min incubation (cf. **Fig 5**).

We further confirmed the direct effect of mouth rinses on SARS-CoV-2 viral infectivity using replication competent viruses expressing mNeonGreen. Similar to the results using pseudotyped viruses expressing spike proteins, 50% (v/v) Listerine, 50% (v/v) CHG, 5% (v/v) Povidone-Iodine, and 5% (v/v) Colgate Peroxyl significantly blocked viral infectivity; 0.5% (v/v) povidone-iodine was not active against the virus, whereas 0.5% (v/v) Colgate Peroxyl had moderate antiviral activity; 5% (v/v) Listerine and 5% (v/v) CHG had a moderate anti-viral effect (**Fig 6A,B**). In contrast to infected cells with exposure to highly diluted povidone-iodine and Colgate Peroxyl during the infection leading to cell death (**Fig 3**), there was no apparent cell death in cells infected by viruses after the removal of mouth rinses by centrifugation. Taken together, Listerine and CHG may be better mouth rinse products for SARS-CoV-2 prevention. Highly diluted Povidone-Iodine and Colgate Peroxyl significantly inactivated viruses but their antiviral effects were associated with severe cytotoxicity.

**Fig. 6.**
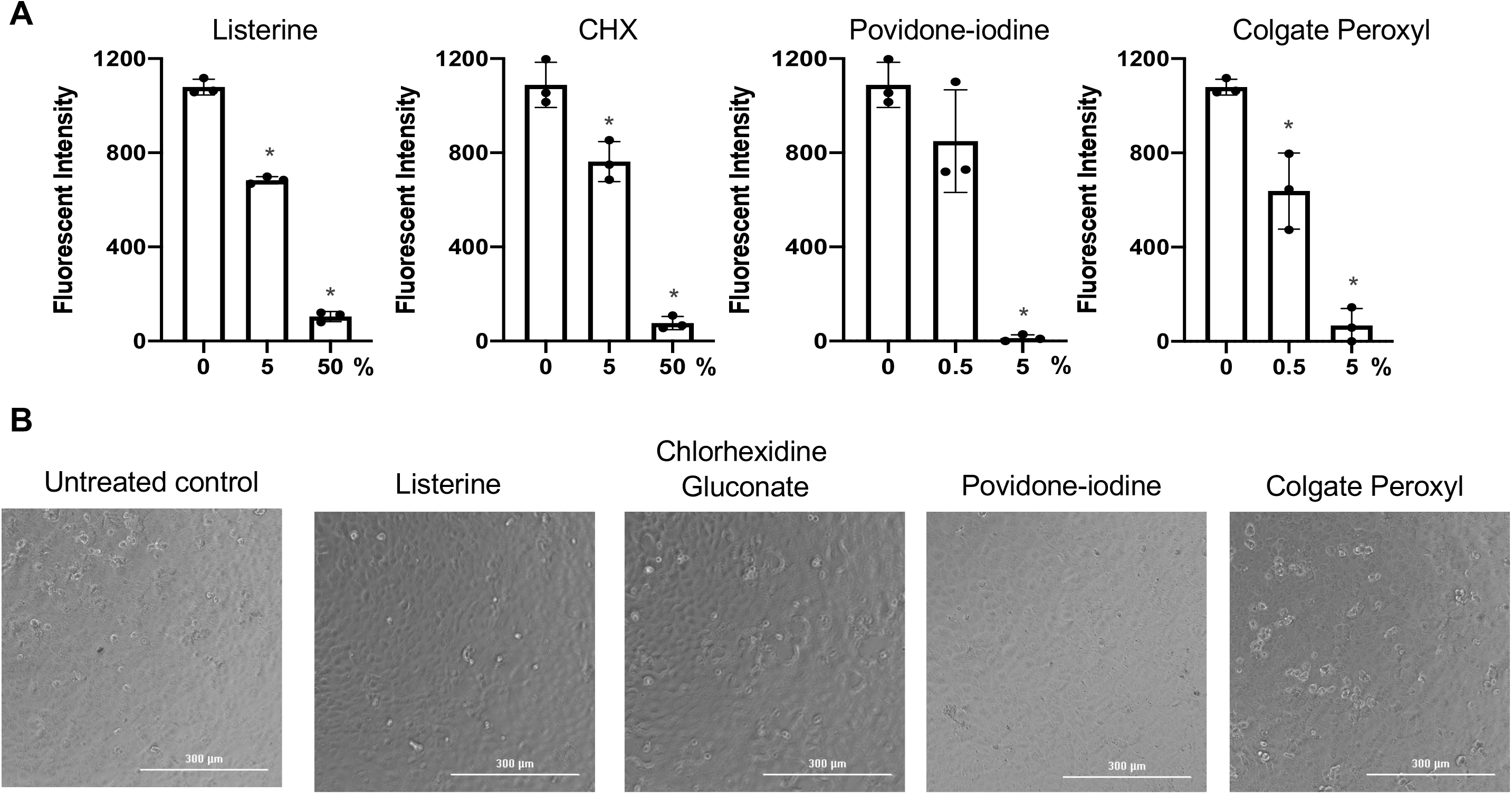
The effect of mouth rinses on replication competent SARS-CoV-2 viruses. (A) Mouth rinses and povidone-iodine at indicated dilutions were added to replication competent SARS-CoV-2 expressing mNeonGreen (1×10^6^ plaque forming units). Treated and untreated viruses were immediately pelleted by centrifugation, and the supernatant was aspirated. Virus pellets were resuspended in Fluorobrite medium with 2% FBS, and added to Vero cells. Viral infection was determined by measuring fluorescence intensity at 24 h post-infection (**Fig 6A**), and images of infected cells with or without mouth rinse treatment were also acquired (**Fig 6B**). Differences between mouth rinse- and medium control (0%)-treated viruses were compared; \**p* < 0.05. Data are means ± SD, and are representative of two independent experiments.

## Discussion

Unlike SARS-CoV and Middle East Respiratory Syndrome (MERS)-CoV, which caused thousands of cases and 700-800 deaths, SARS-CoV-2 appears to be more highly transmissible. The SARS-CoV-2 virus is detectable in saliva from infected individuals without symptoms or with mild symptoms, suggesting that a strategy of suppressing the viral load in the oral cavity may reduce viral spread. Previous studies were conducted to assess the antiseptic effect of Povidone-Iodine on several respiratory viruses, including SARS-CoV, MERS-CoV, and H1N1, using virus-mediated cytopathic effects (Eggers et al. 2018); however, they did not exclude possible antiseptic-associated cytotoxicity. Similarly, published data on the effect of mouth rinses on SARS-CoV-2 infection did not distinguish the impact of mouth rinses on cell viability in efforts to determine the direct effects of mouth rinses on viral infection (Anderson et al. 2020; Bidra et al. 2020; Meister et al. 2020). The apparent effective dosing of the antiseptic rinse on viral infectivity can be misleading when the putative anti-viral effect is accompanied by a cytotoxic effect. Our experiments were designed to discriminate between cytotoxic effects of the mouth rinses and effects of the rinses on the infectivity of the virus. Indeed, we found that anti-viral effects of highly diluted povidone-iodine and Colgate Peroxyl were the consequence of cytotoxicity when the agents were present during a 24 h infection assay. Our results warrant concerns regarding reliability of findings in previous studies in which infected cells were exposed to mouth rinses and antiseptics for extended times.

We found that all mouth rinses tested (all products diluted 1:1 with culture medium, 50% v/v) had cytotoxic effects on cells. We found the cytotoxicity of Colgate Peroxyl > povidone-iodine > CHG > Listerine. Similar trends were observed in both HeLa-hACE2 and oral epithelial cells. Mouth rinse-induced cytotoxicity was more pronounced in cells with 2h incubation than with 20 sec incubation. When CHG was present during a 2-day infection period, 1.5 and 3% (v/v) CHG suppressed SARS-CoV-2 infection by nearly 99% (**Fig 4**). However, 1.5% (v/v) CHG was less potent when the mouth rinse was only present during viral attachment. Importantly, when assessing the effect of CHG on the viruses after removal of mouth rinse during the infection, 5% (v/v) CHG had only a moderate effect, reducing infection by 35-55%. Similarly, potent “anti-viral” effects of 0.1% (v/v) povidone-iodine and 0.05% (v/v) Colgate Peroxyl that were observed when antiseptics were present during infection, were found by the cell image analysis to be due to antiseptic-associated cytotoxicity (**Fig 3**). In fact, we found that 0.5% (v/v) povidone-iodine had little effect on either replication competent or pseudotyped viruses if the povidone-iodine was removed from the virus before infection (**Figs 5 and 6**). Our results show the importance of considering potential cytotoxicity of putative antiviral agents when assessing their anti-viral activities. Despite our finding that commercially available mouthwashes had some degree of cytotoxicity, these formulations are well tolerated in clinical use. The ability to determine the antiviral effects of these mouth washes independent of their cytotoxicity is important for translating these laboratory results into clinical studies.

Both 5% (v/v) Colgate Peroxyl and 5% (v/v) povidone-iodine inactivated virus effectively, whereas 50% Listerine and 50% CHG were required to inactivate the virus. We found that pre-incubation with mouth rinses did not significantly alter the anti-viral profile, indicating that the anti-viral activity occurs rapidly on contact. The determination of the anti-viral activity of diluted mouthwash is important since the salivary flow in the oral and pharyngeal cavities will dilute the activity following application. Thus, assessing the effects of highly dilute mouth rinses over time may help establish the frequency of rinsing necessary for optimal clinical benefits.

The differential anti-viral and cytotoxicity profiles of these mouth rinses suggest that their anti-viral mechanisms are not all the same. The underlying mechanism of anti-viral activity of mouth rinses and their active ingredients remains to be determined. The active compounds in mouth rinses may block infection by altering/disrupting viral envelopes (membranes) and viral proteins. For example, the key ingredient in Colgate Peroxyl is hydrogen peroxide, which is known to increase cell membrane permeability and cause DNA damage (THOMSON 1928; Ward et al. 1985). Hydrogen peroxide may inactivate viruses through lipid oxidation and/or nucleic acid damage. Povidone-iodine blocks influenza A virus by acting on the viral glycoproteins, hemagglutinin and neuraminidase, resulting in inhibition of binding of virus to cells (Sriwilaijaroen et al. 2009). Chlorhexidine, a cationic molecule, reduces infectivity of enveloped viruses, such as human immunodeficiency virus (HIV) and respiratory syncytial virus, and non-enveloped viruses, such as rotavirus and hepatitis A virus (WHO 2009), suggesting that its action is not simply mediated through membrane disruption. Listerine, an essential oil-based mouth rinse, has been shown to inhibit infection by HIV and herpes simplex virus-1 (Baqui et al. 2001), and may reduce the infectivity of viruses through altering hydrophobicity of viral glycoproteins necessary for viral attachment.

In conclusion, all mouth rinses tested inactivated replication competent SARS-CoV-2 viruses and pseudotyped viruses expressing spike proteins. The cytotoxic effects of mouth rinses should be considered when assessing their antiviral activities. Since diluted Listerine and CHG exhibited no cytotoxic effects, these products may be good candidates to reduce virus spread. Studies of antiviral effects of mouth rinses are needed for determining their clinical efficacy in reducing virus spread, particularly in asymptomatic individuals.

## Acknowledgements

We thank Eric C. B. Milner for editing the manuscript. This work was supported by NIH grant NIH R01AI36948 to T.L.C.

## Author contributions

Xu, C: contributed to acquisition, analysis, and interpretation, drafted and critically revised the manuscript.

Wang, A: contributed to acquisition and analysis, and critically revised the manuscript.

Hoskin, E: contributed to conception, interpretation, and critically revised the manuscript.

Cugini, C: contributed to conception, interpretation, and critically revised the manuscript.

Markowitz, K: contributed to conception, interpretation, and critically revised the manuscript.

Chang, T: contributed to design, acquisition, analysis, and interpretation, drafted and critically revised the manuscript.

Fine, D: contributed to conception and design, analysis and interpretation, and critically revised the manuscript.

All authors gave their final approval and agree to be held accountable for all aspects of the work ensuring integrity and accuracy.

---

**Fig. S1.**
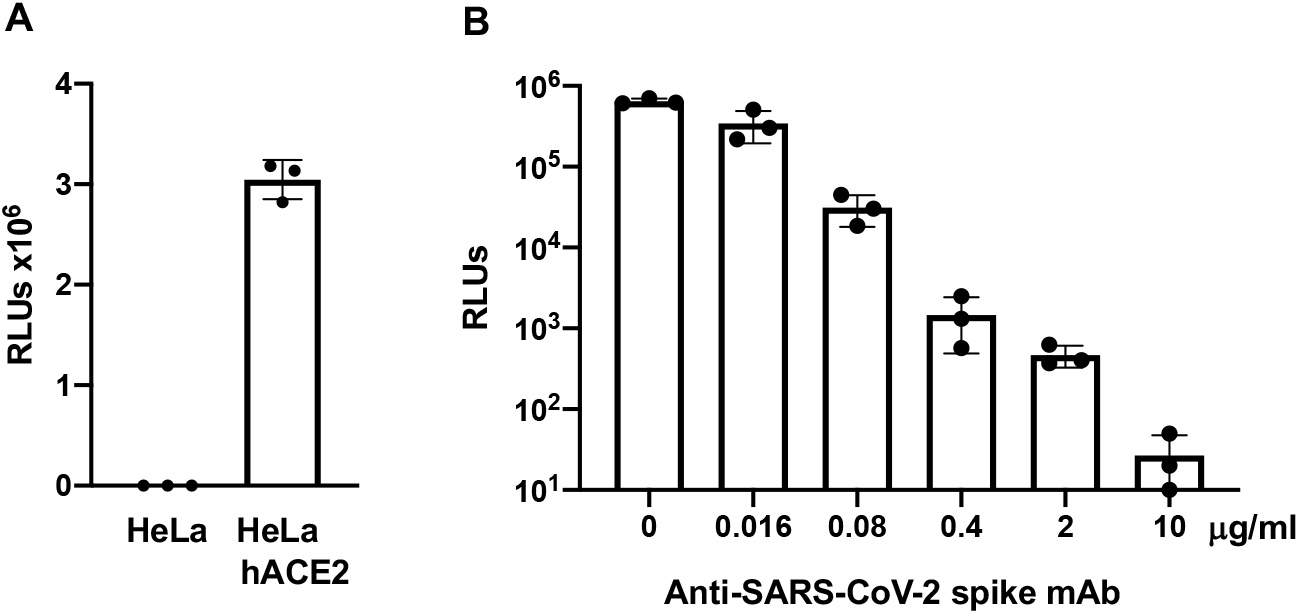
HIV pseudotyped virus expressing SARS-CoV-2 spike proteins. (A) Parental HeLa cells or HeLa cells overexpressing hACE2 were infected with HIV pseudotyped SARS-CoV-2 (~20 ng p24 per well; 48-well plate) for 1 h. Cells were washed, and then cultured for 3 days before measurement of luciferase activity in infected cells. The results show that the infection by pseudotyped luciferase virus expressing SARS-CoV2 spike proteins was hACE2-dependent. (B) Pseudotyped SARS-CoV-2 virus was incubated with different concentrations of monoclonal antibody against SARS-CoV-2 spike proteins for 1 h before infection of HeLa-hACE2 cells. Luciferase activity was measured at 48 h post-infection. The result confirmed that anti-spike protein antibody blocked infection by pseudotyped virus indicating that infection was mediated by the spike proteins.

**Fig. S2.**
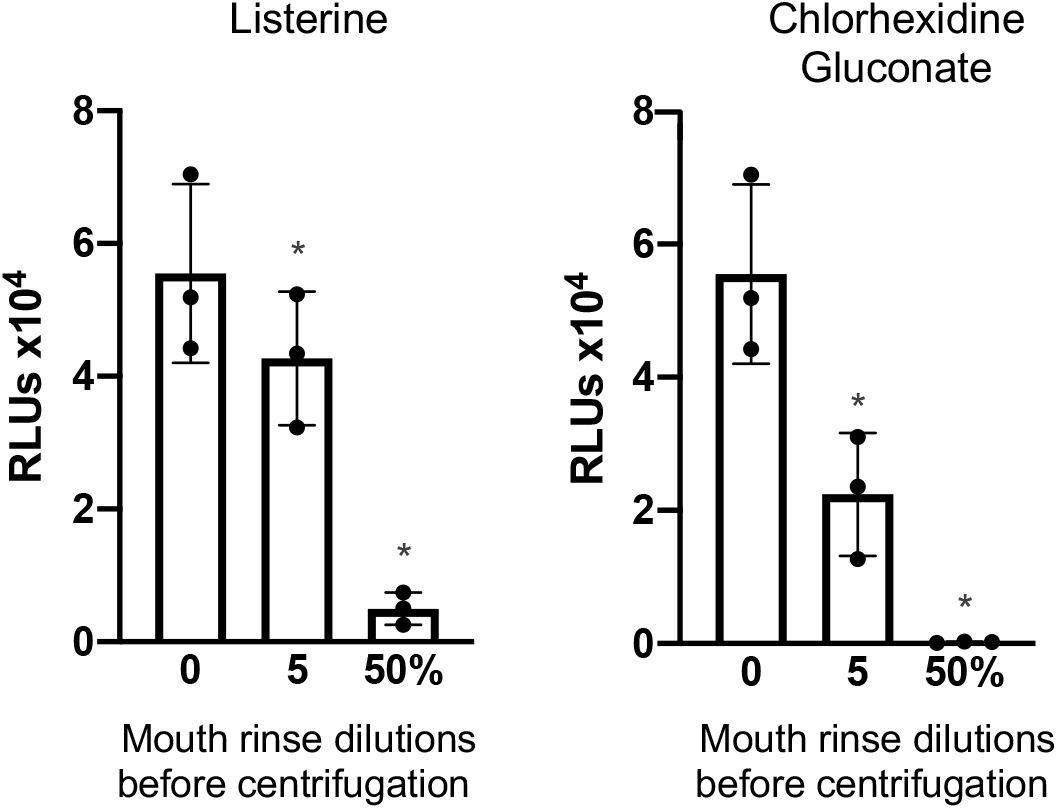
Pre-incubation was not required for the inhibitory effect of mouth rinse on viruses. Mouth rinses were added to pseudotyped SARS-CoV-2 viruses, mixed, and immediately centrifuged. Supernatants were aspirated, virus pellets were resuspended in medium, and added to HeLa-hACE2 cells. Infection was determined by measuring luciferase activity on post-infection day 2. Data are mean ± SD. Differences between mouth rinse-treated viruses and medium control (0%) were compared; \**p* < 0.05. The anti-viral profiles of Listerine and CHG were comparable to the results with pre-incubation of viruses with mouth rinses for 30 min shown in Fig 5, indicating that pre-incubation of viruses with mouth rinses was not required to inactivate the viruses.

## References

Anderson DE, Sivalingam V, Kang AEZ, Ananthanarayanan A, Arumugam H, Jenkins TM, Hadjiat Y, Eggers M. 2020. Povidone-iodine demonstrates rapid in vitro virucidal activity against sars-cov-2, the virus causing covid-19 disease. Infect Dis Ther. 9(3):669–675.

Baqui A, Kelley J, Jabra-Rizk MA, DePaola L, Falkler W, Meiller T. 2001. In vitro effect of oral antiseptics on human immunodeficiency virus-1 and herpes simplex virus type 1. Journal of Clinical Periodontology. 28:610–616.

Bidra AS, Pelletier JS, Westover JB, Frank S, Brown SM, Tessema B. 2020. Rapid in-vitro inactivation of severe acute respiratory syndrome coronavirus 2 (sars-cov-2) using povidone-iodine oral antiseptic rinse. J Prosthodont. 29(6):529–533.

Eggers M, Koburger-Janssen T, Eickmann M, Zorn J. 2018. In vitro bactericidal and virucidal efficacy of povidone-iodine gargle/mouthwash against respiratory and oral tract pathogens. Infect Dis Ther. 7(2):249–259.

Fine DH, Furgang D, Korik I, Olshan A, Barnett ML, Vincent JW. 1993. Reduction of viable bacteria in dental aerosols by preprocedural rinsing with an antiseptic mouthrinse. Am J Dent. 6(5):219–221.

Fine DH, Korik I, Furgang D, Myers R, Olshan A, Barnett ML, Vincent J. 1996. Assessing pre-procedural subgingival irrigation and rinsing with an antiseptic mouthrinse to reduce bacteremia. J Am Dent Assoc. 127(5):641–642, 645–646.

Holmes M, Zhang F, Bieniasz PD. 2015. Single-cell and single-cycle analysis of hiv-1 replication. PLoS Pathog. 11(6):e1004961.

Koletsi D, Belibasakis GN, Eliades T. 2020. Interventions to reduce aerosolized microbes in dental practice: A systematic review with network meta-analysis of randomized controlled trials. J Dent Res. 22034520943574.

Lee S, Kim T, Lee E, Lee C, Kim H, Rhee H, Park SY, Son H-J, Yu S, Park JW et al. 2020. Clinical course and molecular viral shedding among asymptomatic and symptomatic patients with sars-cov-2 infection in a community treatment center in the republic of korea. JAMA Internal Medicine.

Meiller TF, Silva A, Ferreira SM, Jabra-Rizk MA, Kelley JI, DePaola LG. 2005. Efficacy of listerine antiseptic in reducing viral contamination of saliva. J Clin Periodontol. 32(4):341–346.

Meister TL, Bruggemann Y, Todt D, Conzelmann C, Muller JA, Gross R, Munch J, Krawczyk A, Steinmann J, Steinmann J et al. 2020. Virucidal efficacy of different oral rinses against sars-cov-2. J Infect Dis.

O’Donnell VB, Thomas D, Stanton R, Maillard J-Y, Murphy RC, Jones SA, Humphreys I, Wakelam MJO, Fegan C, Wise MP et al. 2020. Potential role of oral rinses targeting the viral lipid envelope in sars-cov-2 infection. Function.zqaa002.

Park JB, Park NH. 1989. Effect of chlorhexidine on the in vitro and in vivo herpes simplex virus infection. Oral Surg Oral Med Oral Pathol. 67(2):149–153.

Sriwilaijaroen N, Hiramatsu H, Takahashi T, Suzuki T, Ito M, Ito Y, Tashiro M, Suzuki Y. 2009. Mechanisms of the action of povidone-iodine against human and avian influenza a viruses: Its effects on hemagglutination and sialidase activities. Virology journal. 6:124.

Thomson DL. 1928. The effect of hydrogen peroxide on the permeability of the cell. Journal of Experimental Biology. 5(3):252–257.

To KK, Tsang OT, Leung WS, Tam AR, Wu TC, Lung DC, Yip CC, Cai JP, Chan JM, Chik TS et al. 2020. Temporal profiles of viral load in posterior oropharyngeal saliva samples and serum antibody responses during infection by sars-cov-2: An observational cohort study. Lancet Infect Dis. 20(5):565–574.

Ward JF, Blakely WF, Joner EI. 1985. Mammalian cells are not killed by DNA single-strand breaks caused by hydroxyl radicals from hydrogen peroxide. Radiat Res. 103(3):383–392.

Wei L, Lin J, Duan X, Huang W, Lu X, Zhou J, Zong Z. 2020. Asymptomatic covid-19 patients can contaminate their surroundings: An environment sampling study. mSphere. 5(3).

WHO. 2009. Who guidelines on hand hygiene in health care: First global patient safety challenge clean care is safer care. https://wwwpahoorg/col/indexphp?option=com_docman&view=download&alias=264-who-guidelines-on-hand-hygiene&category_slug=publicaciones-ops-oms-colombia&Itemid=688.

WHO. 2020. Transmission of sars-cov-2: Implication for infection prevention precautions. World health organization https://wwwwhoint/news-room/commentaries/detail/transmission-of-sars-cov-2-implications-for-infection-prevention-precautions.

Wu F, Zhao S, Yu B, Chen YM, Wang W, Song ZG, Hu Y, Tao ZW, Tian JH, Pei YY et al. 2020. A new coronavirus associated with human respiratory disease in china. Nature. 579(7798):265–269.

Xie X, Muruato A, Lokugamage KG, Narayanan K, Zhang X, Zou J, Liu J, Schindewolf C, Bopp NE, Aguilar PV et al. 2020. An infectious cdna clone of sars-cov-2. Cell Host Microbe. 27(5):841–848 e843.

